# Development of a high dimensional imaging mass cytometry panel to investigate spatial organization of tissue microenvironment in formalin-fixed archival clinical tissues

**DOI:** 10.1101/2022.05.12.491175

**Authors:** Stian Tornaas, Dimitrios Kleftogiannis, Siren Fromreide, Hilde Ytre-Hauge Smeland, Hans Jørgen Aarstad, Olav Karsten Vintermyr, Lars Andreas Akslen, Daniela Elena Costea, Harsh Nitin Dongre

## Abstract

To decipher the interactions between various components of the tumor microenvironment (TME) and tumor cells in a preserved spatial context, a multiparametric approach is essential. In this pursuit, imaging mass cytometry (IMC) emerges as a valuable tool, capable of concurrently analyzing up to 40 parameters at subcellular resolution. In this study, a set of antibodies was selected to spatially resolve multiple cell types and TME elements, including a comprehensive panel targeted at dissecting the heterogeneity of cancer-associated fibroblasts (CAF), a pivotal TME component. This antibody panel was standardized and optimized using formalin-fixed paraffin-embedded tissue (FFPE) samples from different organs/lesions known to express the markers of interest. The final composition of the antibody panel was determined based on the performance of conjugated antibodies in both immunohistochemistry (IHC) and IMC. Tissue images were segmented employing the Steinbock framework. Unsupervised clustering of single-cell data was carried out using a bioinformatics pipeline developed in R program. This paper provides a detailed description of the staining procedure and analysis workflow. Subsequently, the panel underwent validation on clinical FFPE samples from head and neck squamous cell carcinoma (HNSCC). The panel and bioinformatics pipeline established here proved to be robust in characterizing different TME components of HNSCC while maintaining a high degree of spatial detail. The platform we describe shows promise for understanding the clinical implications of TMA heterogeneity in large patient cohorts with FFPE tissues available in diagnostic biobanks worldwide.

## Introduction

Development and progression of cancer is a complex process governed not only by transformed tumor cells but also supported by many types of surrounding non-transformed cells. These surrounding cells include fibroblasts, pericytes, cells related to blood and vascular system, fat, and nerve cells, as well as a myriad of immune cells such as T and B cells, natural killer (NK) cells, myeloid-derived suppressor cells and macrophages [1, 2]. These cells, along with soluble growth factors and extracellular matrix (ECM), comprise the tumor microenvironment (TME) [3]. Fibroblasts within the TME are called cancer-associated fibroblasts (CAFs) and, as recently shown, represent a heterogenous population of cells [2, 4, 5]. CAFs have been proven to support tumor cell proliferation and migration, angiogenesis, immunomodulation and chemoresistance through secretion of different growth factors, chemokines, cytokines or by their own motile phenotype [6-9]. Due to the long-held notion that CAFs support tumor progression, attempts have been made to deplete CAFs in tumors, albeit with poor outcomes [10]. This is owing to their high plasticity and heterogeneity with both pro- and anti-tumorigenic effects [4, 11]. To date there is no consensus on CAF-specific markers, and the majority of studies define CAFs by a combination of spatial location, morphology and lack of expression of linage markers for epithelial cells, leukocytes and endothelial cells [11]. In addition to CAFs, ECM has shown to actively participate in tumorigenic processes by regulating migration and activation of tumor associated immune cells [1]. More comprehensive analysis of CAF and immune cell composition, their tissue spatial distribution and colocalization with ECM components is crucial for understanding both pro- and anti-tumoral mechanisms. This will help to comprehend better the crosstalk between different components of TME and their role in patient prognosis and therapy response.

A multiparametric approach is required to obtain an integrative picture of TME that captures spatial information [12, 13]. Imaging mass cytometry (IMC) allows in-situ characterization of up to 40 markers using metal tagged antibodies which is ideal for characterization of CAF subsets and their crosstalk with TME and tumor cells. Cyclic immunofluorescence has been employed to obtain highly multiplexed images (up to 60 markers) [14] but suffers from spectral overlap, tissue autofluorescence and tissue degradation due to repeated rounds of labeling-stripping-acquisition [14]. The use of metal tagged antibodies in IMC overcomes these issues and allows simultaneous detection of several markers. The technology has been used previously to increase the understanding of several types of cancer [12, 15-17]. Nevertheless, the use of this state-of-the-art technology requires thorough design and validation of markers for optimal performance.

This study describes the development and optimization of a panel of antibodies (AB) that allows effective detection and extensive characterization of CAFs and other TME components in formalin fixed paraffin embedded (FFPE) archival clinical tissues. The developed panel targets fibroblast markers, epithelial and structural markers as well as markers for immune (NK cells, macrophages, T helper cells, cytotoxic T cells, T-regulatory cells, and B cells) and vascular (blood & lymph endothelial cells, pericytes) cells. The panel incorporates several new IMC markers that have been validated using IHC, before and after in-house metal conjugation. In addition, markers/clones that were metal conjugated but failed quality control during IMC acquisition, have also been presented. The panel was used to visualize CAF and TME elements in clinical samples of Head and neck squamous cell carcinoma (HNSCC). Although, the developed AB panel was validated on HNSCC samples, it can be applied to various types of cancer and other pathological conditions such as fibrosis to increase our knowledge about functional roles of fibroblast subpopulations. The main experimental, image pre-processing and post-processing steps used here can also serve as a paradigm for developing panels for other types of cells/pathways both in cancer and in other diseases and it offers the advantage of being usable for analysis of large cohorts of patients with FFPE clinical samples available in diagnostic biobanks.

## Results

### Development, optimization and validation of the antibody panel

To develop a panel for the detection and characterization of CAFs and other TME components via IMC, several (>40) markers/clones were tested (Table 1). All markers included in the study underwent extensive quality control steps with the performance assessed by immunohistochemistry (IHC) both before and after metal conjugation, and then verified by IMC on the cores of a control tissue microarray (cTMA) constructed from tissues/lesions with known expression of markers (Table S1). After metal conjugation, all ABs underwent testing for protocol standardization using different antigen retrieval procedures, primary AB incubation times and temperatures (1hour at RT and overnight at 4ºC). Seventeen ABs were purchased pre-conjugated and except PD-1 (EPR4877), PD-L1 (SP142) and Arginase-1 (D4E3M, Standard Biotools), worked optimally with IHC antigen retrieval buffer pH 9 as suggested by the manufacturer (Figure S1). The rest of the ABs were chosen based on their capacity to optimally work with pH 9 buffer. ABs against VEGFR-3 (five different clones MAB3491, C28G5, C28A2, MM0003-7G63 and D-6), NG-2 (EPR22410-145 and E3B3G), NRP-2 (C-9 and HPA039980), pTAK (90C7), pFAK (D20B1), BNIP3 (D7U1T), VE-cadherin (BV13), Tenascin-c (C-9) did not work with antigen retrieval buffer pH 9 and were not metal conjugated for IMC staining (Figure S2). ABs against VE-cadherin (E6N7A), Arginase-1 (D4E3M, Abcam) and VEGFR2 (D5B1) displayed specific IHC staining but after metal conjugation did not give similar signals, hence were excluded from further analysis (Figure S1). ABs that were found to work optimally with the protocol were then titrated using IMC on cTMA cores using two different dilutions (Figure S3). Finally, spillover matrix was generated to account for signal leak from one channel to another as described previously [18]. This was done by creating a slide individually spotted with same concentration of metal tagged ABs that were used in the final panel. The matrix thus generated from the dataset was used to correct for spillover in the samples.

**Table 1.** Overview of all markers used in study, metal tags used with each AB, concentration, and incubation step.

|  | Marker | Supplier | Clone | Metal | In-house conjugation | Dilution | Incubation |  |
| --- | --- | --- | --- | --- | --- | --- | --- | --- |
|  |  |  |  |  |  |  | Overnight 4°C | RT (1 hr) |
| 1 | α-SMA | Standard Biotoools | 1A4 | 141 Pr | No | 1:400 | x | - |
| 2 | α-11 | UiB | Clone 24 | 153 Eu | Yes | 1:1000 | x | - |
| 3 | FAP | CST | E1V9V | 152 Sm | Yes | 1:1000 | x | - |
| 4 | FSP-1/S100A4 | CST | D9F9D | 172 Yb | Yes | 1:2000 | x | - |
| 5 | CD140β | CST | 28E1 | 149 Sm | Yes | 1:200 | x | - |
| 6 | CD90/Thy-1 | CST | D3V8A | 176 Yb | Yes | 1:1000 | x | - |
| 7 | Ki-67 | Standard Biotoools | B56 | 168 Er | No | 1:200 | x | - |
| 8 | Caveolin-1 | CST | D46G3 | 145 Nd | Yes | 1:2000 | - | x |
| 9 | CD16 | Standard Biotoools | EPR16784 | 146 Nd | No | 1:50 | - | x |
| 10 | CD68 | Standard Biotoools | KP1 | 159 Tb | No | 1:50 | x | - |
| 11 | CD163 | Standard Biotoools | EDHu-1 | 147 Sm | No | 1:600 | x | - |
| 12 | CD4 | Standard Biotoools | EPR6855 | 156 Gd | No | 1:200 | x | - |
| 13 | CD8a | Standard Biotoools | CD8/144B | 162 Dy | No | 1:100 | x | - |
| 14 | CD20 | Standard Biotoools | H1 | 161 Dy | No | 1:400 | x | - |
| 15 | FoxP3 | Standard Biotoools | 236A/E7 | 155 Gd | No | 1:50 | x | - |
| 16 | CD31 | Standard Biotoools | EPR3094 | 151 Eu | No | 1:100 | x | - |
| 17 | VEGFR2 | CST | D5B1 | 165 Ho | Yes | 1:3000 | x | - |
| 18 | Podoplanin | CST | LpMab-12 | 143 Nd | Yes | 1:500 | x | - |
| 19 | CD146 | Abcam | EPR3208 | 175 Lu | Yes | 1:2000 | x | - |
| 20 | Collagen type 1 | Standard Biotoools | Polyclonal | 169 Tm | No | 1:600 | x | - |
| 21 | Tenascin C | Merck | BC-24, C-9 | 163 Dy | Yes | 1:1000 | x | - |
| 22 | E-cadherin | Standard Biotoools | 24E10 | 158 Gd | No | 1:100 | x | - |
| 23 | EGFR | Standard Biotoools | D38B1 | 142 Nd | No | 1:100 | x | - |
| 24 | YAP-1 | CST | D8H1X | 144 Nd | Yes | 1:3000 | x | - |
| 25 | Granzyme B | Standard Biotoools | EPR2012 9-217 | 167 Er | No | 1:100 | - | - |
| 26 | Arginase-1 | Standard Biotoools | D4E3M | 164Dy | No | 1:100 | - | - |
| 27 | PD-1 | Standard Biotoools | EPR4877 (2) | 165Ho | No | 1:100 | - | - |
| 28 | Arginase-1 | Abcam | D4E3M | 139La | Yes | 1:100 | - | - |
| 29 | VE-cadherin | CST | E6N7A | 148 Nd | Yes | 1:25 | - | - |
| 30 | NG-2 | CST | EPR2241 0-145; E3B3G | - | Not conjugated | 1:1000 | - | - |
| 31 | PD-L1 | Standard Biotoools | SP142 | 150 Nd | No | 1:50 | - | - |
| 32 | VEGFR3 | CST | MAB349 1, C28G5, C28A2, MM000 3-7G63 and D-6 | - | Not conjugated | 1:50 – 1:1000 | - | - |
| 33 | pTAK | CST | 90C7 | - | Not conjugated | 1:50 – 1:1000 | - | - |
| 34 | pFAK | CST | D20B1 | - | Not conjugated | 1:50 – 1:1000 | - | - |
| 35 | BNIP3 | CST | D7U1T | - | Not conjugated | 1:50 – 1:1000 | - | - |
| 36 | NRP2 | AH diagnostics | C-9 and HPA039 980 | - | Not conjugated | 1:50 – 1:1000 | - | - |

### Exploration of cellular spatial localization and cell segmentation

In total, twenty-four markers were validated to perform well with both IHC and IMC. Of those, fourteen ABs were pre-conjugated (α-SMA, Ki-67, CD16, CD68, CD163, CD4, CD8a, CD20, FOXP3, CD31, collagen-1, E-cadherin, EGFR, and granzyme B), and ten were in-house conjugated (integrin α-11, FAP, FSP-1, CD140β, CD90, caveolin-1, podoplanin, CD146, Tenascin-C and YAP-1) (Table 1). When compared to IHC, most of the ABs demonstrated comparable staining patterns in IMC, although the staining intensity was different in some cases (Figure 1, 2, 3).

**Figure 1.**
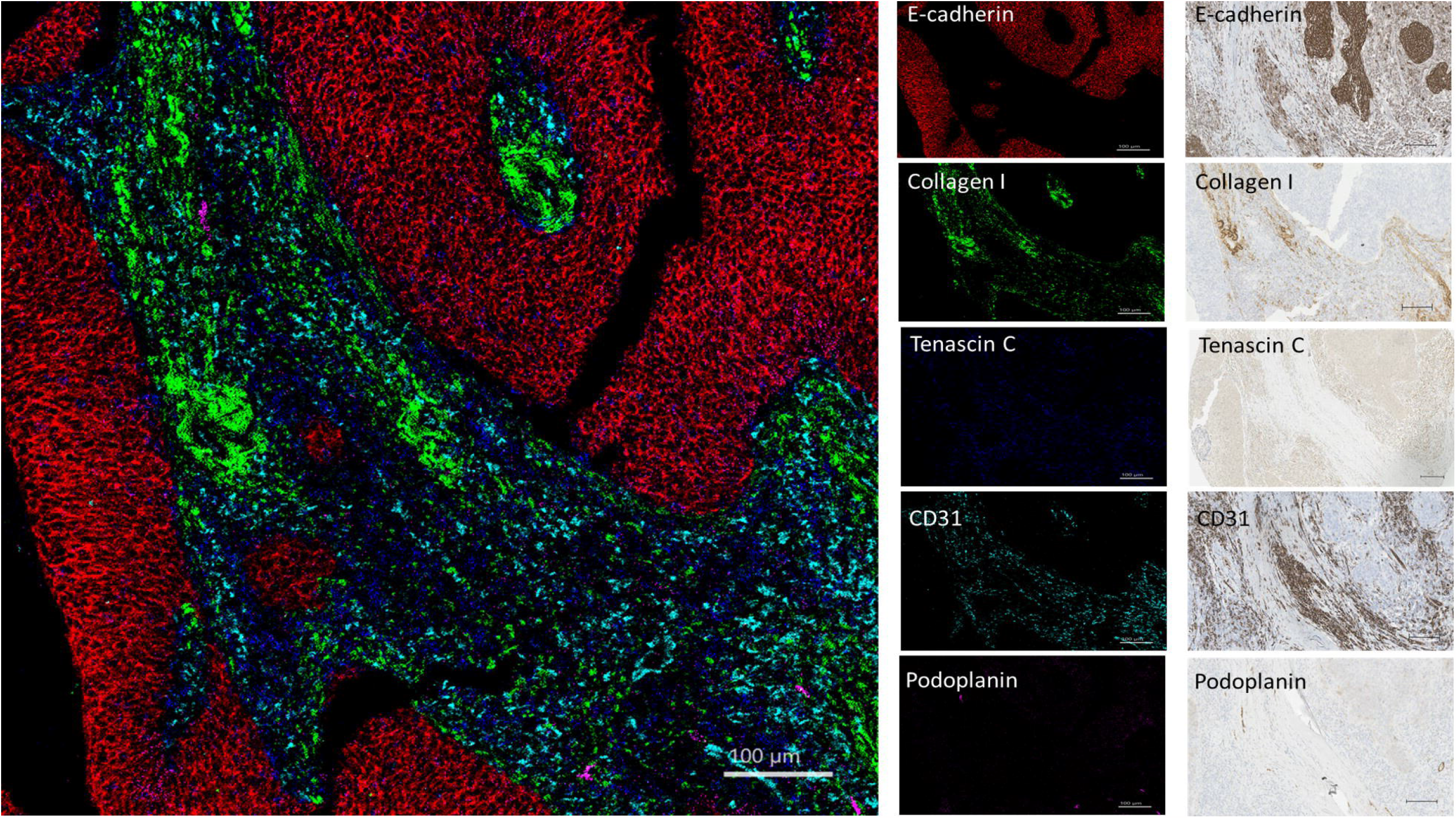
Visualization of structural, stromal, and vascular components of tumor by imaging mass cytometry and immunohistochemistry in an HNSCC core from cTMA. E-cadherin (red) staining the tumor epithelial compartment, collagen-1 (green) and tenascin-C (blue) staining the stroma and CD31 (cyan) and podoplanin (magenta) staining the blood vessels and lymph vessels, respectively. Scale bar-100 μm for IMC and IHC images.

**Figure 2.**
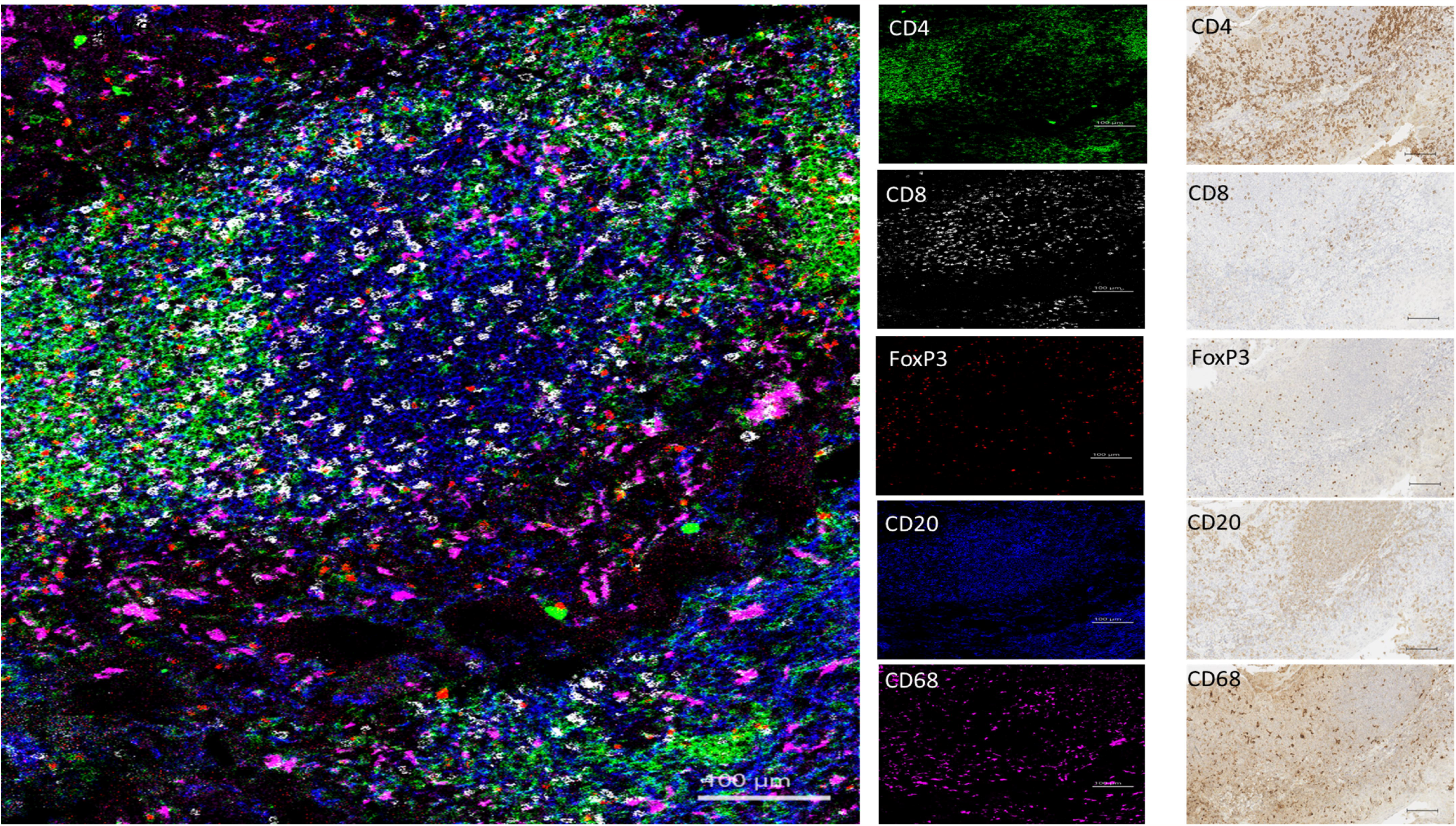
Simultaneous visualization of various immune cell subsets in a tonsil core from cTMA: T-helper cells (green, CD4^+^), cytotoxic T-cells (white, CD8^+^), regulatory T-cells (CD4^+^, FoxP3^+^, red), B-cells (blue, CD20^+^) and macrophages (magenta, CD68^+^). Scale bar-100 μm for IMC and IHC images. FoxP3: forkhead box P3.

**Figure 3.**
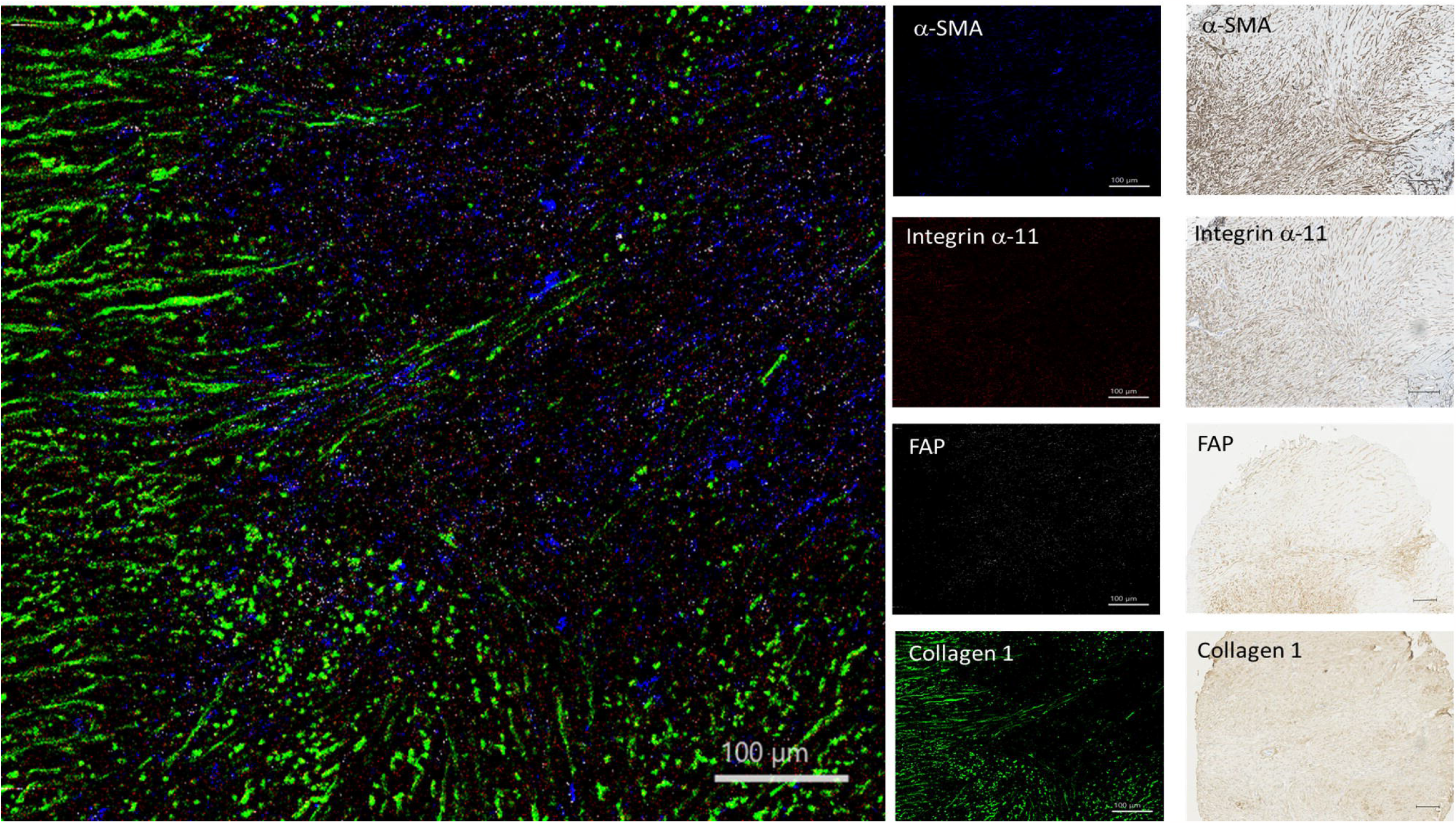
Detection of different CAF markers in osteosarcoma core from cTMA by imaging mass cytometry and immunohistochemistry. CAFs expressing α-SMA (blue), FAP (white), integrin α11 (red) and collagen-1 (green) could be detected. Scale bar-100 μm for IMC and IHC images. α-SMA: alpha-smooth muscle actin, FAP: fibroblast activation protein.

The AB panel described in this study focused on structural markers (as exemplified in the HNSCC core from cTMA, Figure 1), immune cell subsets (tonsil core from cTMA, Figure 2) and fibroblast markers (osteosarcoma core from cTMA, Figure 3). HNSCC tissue core containing structural and vascular compartments was chosen to validate expression of E-cadherin, Collagen-1, Tenascin-C, CD31 and podoplanin. These markers were further used to visualize cells related to epithelial, stromal, and vascular tumor compartments, respectively. E-cadherin was restricted to the epithelial compartment whereas collagen-1 and tenascin-C revealed the stromal compartment of the tumor as observed by both IMC and IHC (Figure 1). In addition, in the same cTMA core most cancer cells showed EGFR expression, whereas Ki-67 depicted proliferating cells in both the cancer and stromal compartment (Supplementary Figure S4). In the tonsil cTMA core, various immune cell types were identified – in a similar way by both IMC and IHC: CD4^+^ (T helper cells), CD8^+^ (cytotoxic T cells), CD20^+^ (B cells), CD4^+^ FoxP3^+^ (T regulatory cells) and CD68^+^ (macrophages) (Figure 2). Moreover, CD16^+^ (natural killer cells) and CD163^+^ (M2 macrophages) could be detected in the HNSCC core of the cTMA (Supplementary Figure S5). The frequencies of Foxp3+, CD20+, CD4+, CD68+, CD163+, CD8+ and Ki67+ cells determined by IMC were qualitatively similar with the matched IHC scores (Supplementary Figure S5). Osteosarcoma tissue core from the cTMA was chosen to validate the presence of the CAF markers: α-SMA, integrin α11, FAP, collagen-1, FSP-1, CD140β, caveolin-1 and CD90 (Figure 3 and Supplementary Figure S6). α-SMA and FAP were observed to be expressed by one subset of cancer cells and integrin α11 and collagen-1 by another in IMC. Similarly comparable staining of FSP-1, CD140β and caveolin-1 was observed between IMC and IHC in the osteosarcoma core of the cTMA (Supplementary Figure S6).

Images from four cores of the cTMA (HNSCC, tonsil, placenta and osteosarcoma) were segmented into single cells using Steinbock computational framework developed by Bodenmiller laboratory [19]. A total of 22,996 cells from all four cores were identified, with HNSCC having 10,088 segmented cells (43.9%), followed by tonsil (36.4 %), placenta (10.5 %) and osteosarcoma (9.2 %). Further, based on prior knowledge of cell type defining markers, segmented cells were clustered. Combined for all cores, 43.96% were immune cells, 22.17% were epithelial cells, 4.36% were endothelial cells, 0.58% were pericytes, 15.76% were fibroblasts and the rest (13.17%) remained unassigned (low expression of all markers) (Figure 4A, B, C). Of note, most of the unassigned cells belonged to the HNSCC core suggesting cell types not explored by our panel. In the tonsil tissue, mainly immune cell subtypes were identified, with a majority of them being CD20+ B cells and their spatial distribution suggesting the presence of a germinal center (Figure 4C). Two-dimensional graphs using the dimensionality reduction algorithm t-distributed stochastic neighbor embedding (t-SNE) were generated to visualize cells from different samples and cell types in each sample. t-SNE plots confirmed presence of different cell lineages with most cells from the tonsil core expressing various immune markers, those from the osteosarcoma core expressing fibroblast markers and those from placenta core expressing both structural and vascular markers. Interestingly, placenta-specific and osteosarcoma-specific clusters separated from both HNSCC and tonsil clusters as well as from each other (Figure 4D, E). Although, a HNSCC-dominant cluster could be seen containing mainly epithelial markers, HNSCC-cells were spread across all the markers, suggesting presence of different cell lineages in the HNSCC tissue.

**Figure 4.**
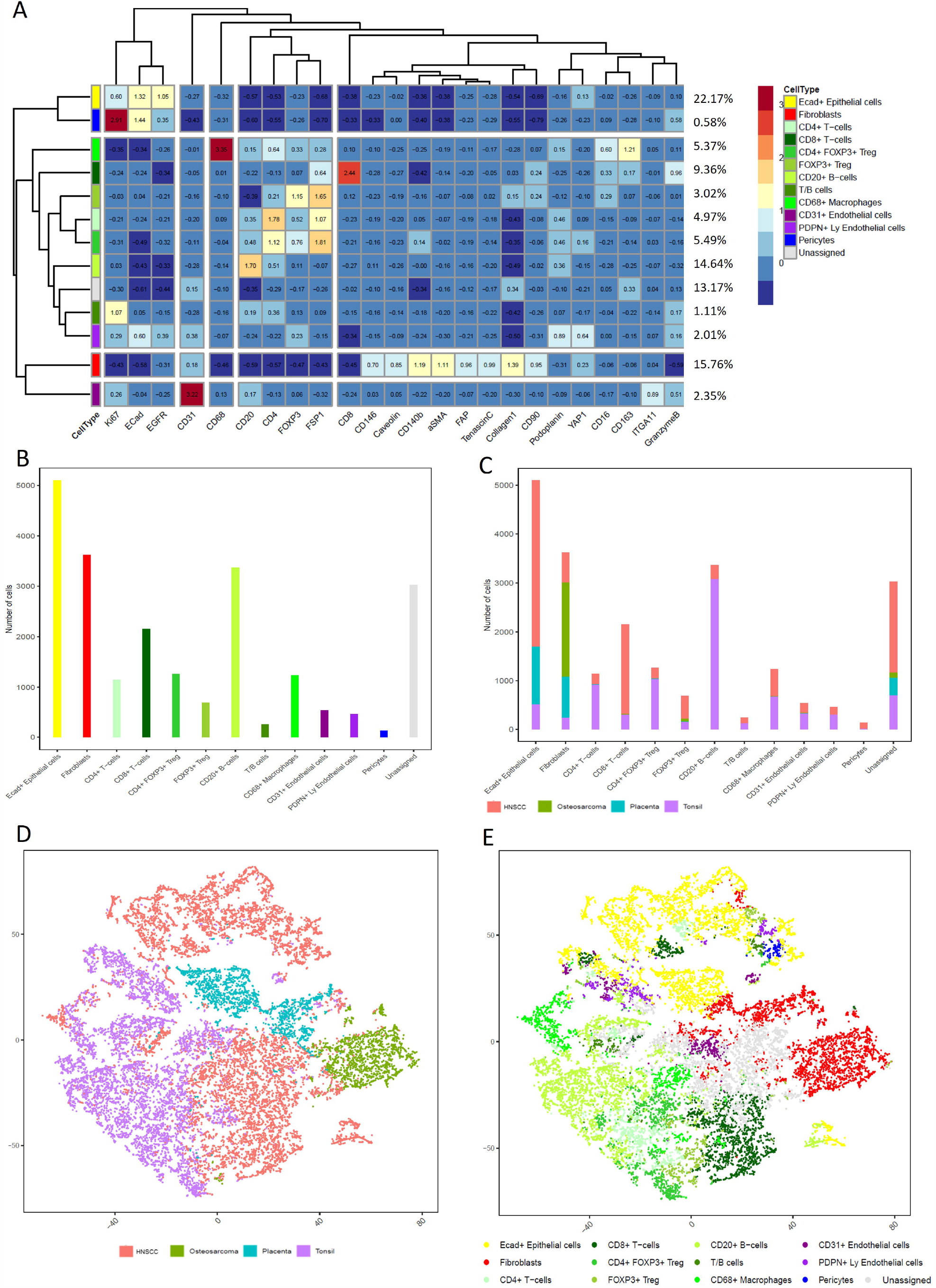
Single-cell clustering using R pipeline. A) Heatmap depicting thirteen major clusters based on semi-supervised clustering. All major cell types could be identified. B) and C) Number of cells and relative percentage in each cluster related to tissue type. D) Distribution of cells according to tissue type visualized after dimensionality reduction using t-SNE. E) Distribution of cells based on marker expression and phenotype in the same t-SNE. T-SNE: t-distributed Stochastic Neighbor Embedding. HNSCC: Head and neck squamous cell carcinoma. FSP-1: fibroblast specific protein-1, α-SMA: alpha-smooth muscle actin, FAP: fibroblast activation protein.

### Unsupervised clustering identified major cell populations in head and neck squamous cell carcinoma clinical samples

The established IMC panel was then used to identify and visualize TME components in a pilot cohort of HNSCC samples (n=5, two tTMA cores per sample, Table S2). The acquired ROI images were segmented into single cells using the described workflow. Single cell segmentation discovered 89,358 cells from the five samples, ranging from 12,396 to 22,230 cells per tumor (median 18,196 cells/tumor). Based on unsupervised clustering of single cell expression data, 115 cell clusters were identified. Clusters having similar expression patterns were then merged resulting in fourteen meta clusters (Figure 5A). All major cell types could be identified with E-cadherin+ epithelial cells accounting for around 30% of cells and were the most abundant cell type (Supplementary figure S7). On the other hand, CD146+ pericytes were the least abundant cell type (0.62%). Among the immune component, CD4+ T cells were most abundant (11.72%) followed by CD8+ T cells (10.6%) and CD68+ macrophages (10.25%) (Figure 5B). Of note, cell populations expressing both CD20 and CD4 were also found and defined as T/B cells, consistent with previous findings [17]. t-SNE plots generated from all samples did not reveal global differences among the samples or between different cores from same patient (Supplementary Figure S7). Interestingly, two clusters were negative for other major cell lineage markers and positive for stromal markers. These were defined as CAF-1 and CAF-2. CAF-1 showed high expression of αSMA, CD140β and CD90 whereas CAF-2 were high in FAP and collagen-1 expression. t-SNE plots for different cell types validated findings of two CAF subtypes as the two clusters were spatially distant from each other (Supplementary Figure 7). When the different cell types were mapped on tumor cores, CAF-2 were found to be closely located around cancer cells whereas CAF-1 subtype was found closer to immune cells (Figure 5 C, D).

**Figure 5.**
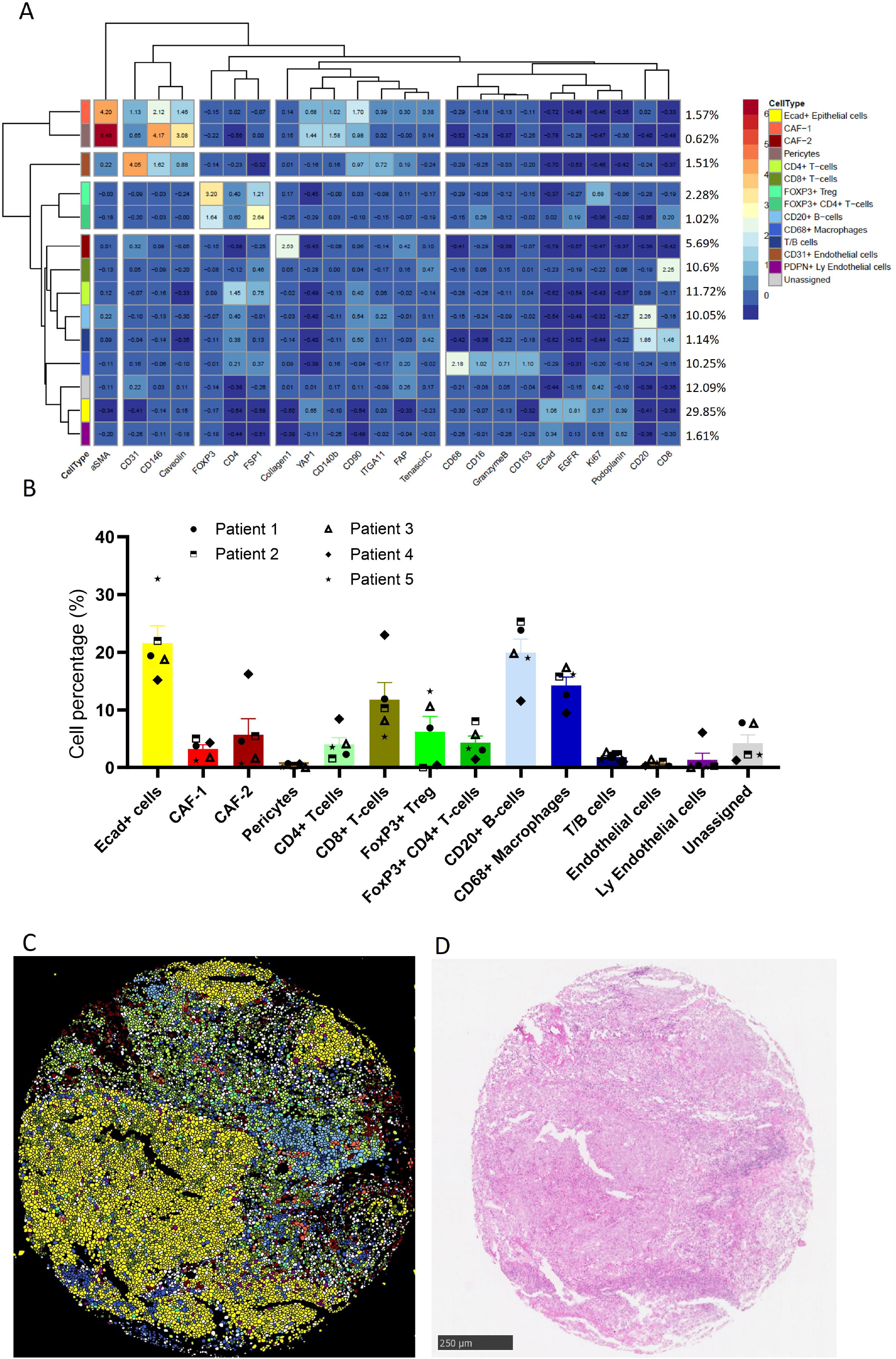
Visualization of different cell types in HNSCC tissue. A) Heatmap identifying various cell types in HNSCC samples (n=5, two cores each). B) percentage abundance of various cell types identified by semi-supervised clustering. C) and D) Visual comparison of segmented cells using in-house bioinformatics pipeline and HE stained core. HNSCC: Head and neck squamous cell carcinoma. FSP-1: fibroblast specific protein-1, α-SMA: alpha-smooth muscle actin, FAP: fibroblast activation protein.

## Discussion

IHC is a widely used technique in biomarker discovery, and a common diagnostic tool in clinical practice [20, 21]. However, IHC is limited by the number of ABs that can be used simultaneously and thereby limiting the information obtained. Cancer is daunting in its breadth and heterogeneity due to genetics, differences in cell and tissue biology, response to therapy and pathology. With the development of immunotherapy, more powerful experimental and precise tools are required to provide a “snapshot” of the tumor along with its microenvironment. IMC is one such technique that has the possibility to use up to 40 different ABs simultaneously [16, 22]. IMC is a low throughput technique generating high dimensional “big data” applicable on both frozen and FFPE tissue [23]. A major prerequisite for any technique to work on archival FFPE tissues is the need of antigen retrieval which greatly affects antibody performance. A multiplex protocol like the one used in IMC, therefore, requires thorough validation. In this study, we illustrate the development of an antibody panel optimized to define CAFs, characterize CAF heterogeneity and other TME components and we validate it on clinical samples of head and neck squamous cell carcinoma. This is a huge improvement on simultaneous detection of protein expression in tissues compared to IHC making IMC better suited at capturing the vast complexities of tumor and its microenvironment.

A better understanding of TME and its *in situ* characterization of spatial localization of different cell types is imperative for identifying clinically relevant single-cell signatures for improved patient care. Amongst the cells found in the TME, CAFs play a dual role, being shown to both enhance tumor cell proliferation, invasion and metastasis but also prevent cancer cell invasion [24]. This dual role has been explained by variety of markers used to define CAFs and lack of a general definition [11]. Eight distinct CAF markers αSMA, collagen-1, CAV-1, CD140β, FAP, integrin α11, FSP-1 and CD90 were optimized during this study and validated on five HNSCC tumor samples. In addition, CAFs are known to mediate response to immune checkpoint inhibitors (ICI) therapy [25] and hence it is essential to decipher the CAF heterogeneity and their crosstalk with immune cells to identify therapeutic targets for future combinatorial trials. Our IMC panel represents a comprehensive set of targets for spatial characterization of CAFs and their interactions with immune cells and other TME components. It includes eight markers to identify different immune cell subsets and their spatial localization to CAFs and tumor cells. Moreover, our panel also includes markers to identify vascular components, both blood and lymphatic vessels. Our panel and subsequent downstream analysis paradigm could aptly differentiate between different tumor compartments.

The main challenge for optimal IMC panel design is to use the same antigen retrieval conditions. This is further complicated by the fact that commercially available pre-conjugated IMC antibodies are validated to work with antigen retrieval pH 9 buffer [26]. In our panel, 35 different ABs were tested out of which seventeen were purchased pre-conjugated. For the rest, thorough literature search was performed to obtain ABs that were compatible with antigen retrieval buffer pH 9 and were available in carrier-free format for metal conjugation. Eighteen ABs and clones met these criteria and testing was done on cTMA cores first with IHC. The cTMA cores chosen to exemplify our optimization work were selected based on the predominance of one TME compartment: HNSCC for its structural component, osteosarcoma for stromal component, placenta for vascular component, and tonsil for its immune-related component. When satisfactory results were obtained, ABs were metal conjugated and tested again with IHC first and then finally titrated with IMC. ABs against VE-cadherin (E6N7A), Arginase-1 (D4E3M, Abcam) and PD-1 (EPR4877) did not work after metal conjugation. This could be explained by the fact that conformational changes or destruction of antigen binding sites during metal conjugation step might have occurred [27]. Lastly, pre-conjugated ABs purchased from supplier, even though pathologist verified needs to be validated again.

A typical IMC pipeline includes image acquisition, single cell segmentation, single cell data transformation/normalization and phenotypic annotation of single cells typically via unsupervised clustering [28]. In our study, single cell segmentation was done with Steinbock framework including DeepCell developed by Windhager et al [19]. Segmentation of cells using Steinbock is an automated process where markers representing nucleus and cytoplasm are selected. The segmentation process is performed using deep convolutional neural networks, which are considered state-of-the-art models for image analysis[19]. This is a huge advantage compared to manual segmentation using Ilastik and CellProfiler, because it greatly reduces the time needed when working with huge data sets [28]. Further, single-cell data normalization and phenotypic annotation of single cells was done by using Histology Topography Cytometry Analysis Toolbox (HistoCAT). With HistoCAT it is also possible to cluster cells based on marker expression, obtaining single-cell information and discover cell neighborhoods, however the program has significant drawbacks [19]. It does not allow spatial visualization of cells in tissues and does not scale well as the image volume increases [29, 30]. Another alternative that could be useful for downstream analysis of single-cell data is ImaCytE, which is very easy to use and gives the possibility to cluster data, generate heatmaps, cohort comparison and easily visualize cells in tissues and interactions between them [31]. Thanks to the recent advances in single-cell research, a variety of pipelines are available utilizing different analysis platforms (python, R, MATLAB etc..). In addition, various computational methods can be used to extract and perform spatial analysis of IMC data such as “analysis of cancer tissue microenvironment” (LOCATOR) approach [32] or monkeybread [33]. We have used functionalities from Cytomapper package and adopted them in R to investigate the spatial location of single-cells and visualize the tissue structures [34]. Nevertheless, we acknowledge that our study was basic in terms of bioinformatic analysis and was intended to mainly demonstrate the potential of established antibody panel. Nonetheless, it is important to have thorough quality control steps for designing AB panel and downstream analysis as the resolution and the detail at which groups of phenotypically similar cells clustered together depends strongly on the AB panel, the tissues of interest, focus of the studied cell types and the tuning of clustering algorithms.

Despite intense research on the role of CAFs in tumor initiation, progression and resistance to therapy the mechanisms are yet to be fully understood [35]. Unveiling CAF heterogeneity and the functions of various subclasses might improve personalized medicine approach aiming to deplete CAFs that are pro-tumorigenic and boost CAFs that are anti-tumorigenic. To our understanding, there are no established IMC panels having a focus on CAF heterogeneity and, therefore, this work will support the future work of understanding these cell types, their interactions with other TME components and the implications for their heterogeneity on clinical outcome.

## Acknowledgments

The authors would like to acknowledge the Flow Cytometry Core Facility, Department of Clinical Science, University of Bergen for their assistance in acquisition. This work was supported by the Research Council of Norway through its Centers of Excellence funding scheme, (Grant No. 22325) and The Western Norway Regional Health Authority (Helse Vest Grant No. 912260/2019).

## Author Contributions

Conceptualization, ST., DK., DEC. and HD.; data curation, ST, DK and HD.; formal analysis, ST, DK and HD.; funding acquisition, LA, OKV, HJA and DEC.; investigation, ST, DK, SF, HYHS, DEC and HD.; methodology, ST, DK, SF, HYHS, HJA, OKV, LA, DEC and HD.; project administration, DEC and HD.; supervision, DK, LA, DEC and HD.; validation, DK, ST, SF and HD.; visualization, ST, DK and HD.; writing—original draft, ST.; writing—review & editing, ST, DK, SF, HYHS, HJA, OKV, LA, DEC and HD. All authors have read and agreed to the published version of the manuscript

## Material and methods

### Patient material

Archival FFPE blocks were used to prepare both the “control” and the ‘test’ tissue micro arrays (cTMA and tTMA, respectively). The FFPE tissues were chosen based on documented positive expression of markers in different types of tissues. The cTMA samples used for testing as positive controls are listed in Supplementary Table S1. Further, five patient samples with primary diagnosis of Head and neck squamous cell carcinoma (HNSCC) were included in the ‘test’ cohort. All patients were undergoing treatment at the Haukeland University Hospital, Bergen, Norway. Patient samples were anonymized, and all patient informed consents were collected according to the Declaration of Helsinki. All research was approved by the Regional Committee for Medical and Health Research Ethics (REK vest: 2010/481;2011/125).

### Antibodies and metal conjugation

Different clones of ABs were selected based on available literature and in-house knowledge (Diagnostic Pathology Service, Laboratory Clinic, Haukeland University Hospital, Bergen-Norway) with a particular focus on CAF markers. ABs in carrier-free solution were conjugated to purified metals in the lanthanide series using suppliers’ protocol (Standard Biotools, CA, USA). Conjugated AB were eluted in 30 μL antibody stabilizer solution (Standard Biotools, CA, USA) supplemented with 0.05 %sodium azide (Merck, Darmstadt, Germany) to a total volume of 50 μL stored in low attachment 1 mL Eppendorf tubes.

### Immunohistochemistry

From the TMA blocks, four μm-thin tissue sections were cut and placed on glass slides (Thermo Scientific, J1800AMNZ, MA, USA) and subsequent staining was performed. ABs for IMC were first tested with IHC both before and after metal conjugation to validate each AB staining quality. For this, tissue sections were baked over night at 60ºC and rehydrated with decreasing concentration of alcohol. Antigen retrieval was done using pH 9 (Dako, S2367, CA, USA) solution in a pressure boiler (BioCare medical Model DC2008INTL, CA, USA) at 120ºC for 10 min. Endogenous peroxidase activity was blocked with 3 % hydrogen peroxidase solution (Dako, S2023, CA, USA) for 5 min. Further, sections were washed with tris buffered saline with tween (TBST) for 10 min before unspecific protein binding was blocked using 10 % goat serum (Dako, X0907, CA, USA) in 3 % bovine serum albumin (BSA) (Merck, A3059, Darmstadt, Germany) for 20 min. Sections were then stained with primary AB for either 60 min room temperature (RT) or overnight at 4º C (Table 1). After washing with TBST twice for 5 min each, slides were incubated with secondary horseradish peroxidase (HRP) conjugated (Dako, K8000, CA, USA) AB for 30 min followed by chromogen diaminobenzidine (DAB) (Dako, K8012, CA, USA) staining for 10 min for AB detection. Slides were then counterstained with hematoxylin (Dako, S3301, CA, USA) and dehydrated by increasing concentration of alcohol and xylene before mounting with cover glass using pertex (HistoLab, 00811-EX, Askim, Sweden). Slides were scanned using whole slide digital scanner (Hamamatsu NaNoZoomer-XR, Shizuoka, Japan) and visualized using NDP.view2.

### Protocol for IMC staining and data acquisition

Tissue sections were deparaffinized in fresh xylene twice for 10 min each followed by rehydration in descending grades of fresh ethanol (100 %, 95 %, 70 %) for 5 min in each grade. Slides were then washed with MilliQ water for 5 min before antigen retrieval was done using pressure cooker at 120ºC for 10 min. Samples were then cooled down and first washed with MilliQ water for 10 min and then with MaxPar PBS (Standard Biotools, CA, USA) for 10 min. Hydrophobic barrier was drawn followed by blocking with 3 % bovine serum albumin (BSA) in MaxPar PBS for 45 min at RT in a humid chamber. The blocking solution was shaken off and AB mix was applied and incubated at 4ºC overnight. The next day, slides were washed with MaxPar PBS supplemented with 0.2 % Triton X100 (PBS-T, Merck, Darmstadt, Germany) thrice for 5 min each and then applying next round of AB mix, which were incubated for 1 hour at RT. Slides were washed again with Maxpar PBS-T (where from) thrice for 5 min each followed by incubation with Iridium (Ir) solution (Standard Biotools, CA, USA) for 5 min. Slides were washed thrice with MaxPar PBS 5 min each and then briefly with MilliQ before air-drying the sections.

Images were acquired using Hyperion imaging system (Standard Biotools) and the system was tuned according to manufacturer’s protocol. Regions of interest were selected from four cTMA samples and ten cores from five HNSCC samples, and the areas were laser-ablated at 200 Hz which produces pixel size/resolution of 1 μm^2^. The acquisition was done using CyTOF software v7.0 (Standard Biotools). The visualization and success of producing high-quality images with selected combination of markers was done using MCD Viewer v1.0.560.6 (Standard Biotools).

### Image preprocessing and cell segmentation

Segmentation of raw IMC data was done following the Steinbock framework with Docker container (Deepcell) [19]. Briefly, IMC data was processed by running commands with command prompt (CMD, Windows). A panel.csv file was generated and channels for segmentation (nucleus and cytoplasm/membrane) were chosen. In this study, channels Ir191 and Ir193 were used as nuclear markers, while αSMA, EGFR, CD31, CD4, E-cadherin, CD20 and FSP-1 were used as cytoplasmatic/membrane markers. The end-result was cell masks for each tissue type included in the study. Cell segmentation were quality checked with ImaCytE. For this, mask generated as described above were organized in folders, one for each ROI with image stack ome.tiff files representing all channels from each ROI that were exported from MCD viewer. When segmentation was confirmed with ImaCytE, Steinbock was used to export folder with each individual channel. These folders/images were imported into histoCAT [29] for generation of .csv files that was used for further annotation of single cells into similar clusters in R.

### Spillover correction

Spillover correction of IMC data was done following the protocol established by Bodenmiller laboratory [18]. A super frost glass slide (Thermo Fisher Scientific, 10748721, USA) was dipped in 2 % agarose solution (w/v) and left to dry overnight in a vertical position at RT. The following day, a grid was drawn on top of the solidified agarose as reference. Then, 0.3μL of 0.4 % trypan blue was pipetted to the center of each square and left to dry for 2 hours. This was followed by pipetting 0.3μL of each antibody conjugate on top of trypan blue and air dried overnight at RT. ROI was made for each spot/antibody with an area of 200 pixels x 10 pixels and acquisition was done based on parameters described above.

### Unsupervised clustering of single cells

Single cells from each tissue cores were clustered into groups of phenotypically similar cells based on an unsupervised clustering approach. Initially the FlowSOM algorithm [36] was used to generate 225 (grid of 15x15) group of cells. For high-dimensional clustering of single-cells, 24 markers were used namely: α-SMA, caveolin-1, collagen-1, CD4, CD8a, CD16, CD20, CD31, CD68, CD90/Thy-1, CD140β, CD146, CD163, E-cadherin, EGFR, FAP, FoxP3, FSP-1/S100A4, integrin α-11, Ki-67, podoplanin, tenascin-C, granzyme B, and YAP-1. The resulting clusters were aggregated based on the ConsensusClustering method [37] with Euclidean distance metric and average linkage. To assess the quality of each aggregated meta-clustering solution, the Proportion of Ambiguous Clustering (PAC) metric was estimated [38]. To maintain relatively low-level ambiguity, and in order to detect rare groups of phenotypically similar cells, meta-clusters at a PAC cutoff of 1% were merged. At this PAC level, 115 meta-clusters were obtained. Further, meta-clusters that showed a marker expression typical of endothelial, immune, epithelial cells, and stromal cells were aggregated together. One additional phenotypic subgroup was included to annotate all other single cells that exhibited “unassigned” marker expression profiles. For visualization of high-dimensional single-cell data into the 2D space we utilized the t-SNE [39] dimensionality reduction algorithm. The full set of 24 markers was used as input and the algorithm was run with default parameters (perplexity = 30, theta = 0.5) on all cells (n=22996).

### Bioinformatics analysis workflow and code/data availability

The single cell analysis workflow described here was implemented in R, and the codes are publicly available at https://github.com/StiThor/IMC_data_analysis [40]. The workflow allows users to perform very basic analysis for example to extract single-cell data information from segmented IMC images, cluster phenotypical similar cells, and perform exploratory visualization of single-cells using heatmaps and t-SNE plots. To investigate the spatial location of single-cells, and visualize the tissue structures, functionalities from Cytomapper package were adopted [34], and tissue visualization routines have been provided in the code.

## Supplemental information titles and legends

**Supplementary Table 1**. Overview of tissue types (control tissue micro array) used in cTMA construction

**Supplementary Table 2**. Table S2 Clinical parameters of patients in tTMA

**Supplementary Figure 1**. ABs against PD-1 (EPR4877), PD-L1 (SP142), and Arginase-1 (D4E3M) demonstrated good results with IHC, but with IMC, showed non-specific binding and a different staining profile to IHC. VEGFR2 (D5B1) and VE-cadherin (E6N7A) gave high non-specific background signal in IMC after metal conjugation. Placenta core from cTMA was used to compare findings between IHC and IMC for VE-cadherin and HNSCC core for the rest of ABs. Scale bar 100um.

**Supplementary Figure 2**. Antibodies against VEGFR-3 (MAB3491, C28G5 and MM0003-7G63), NRP-2 (C-9), and VE-cadherin (BV13) did not work with IHC on placenta core using antigen retrieval buffer pH 9 whereas ABs against NG-2 (EPR22410-145), pFAK (D20B1), Tenascin-c (C-9) BNIP3 (D7U1T) and pTAK (90C7) showed no specific IHC staining on HNSCC core under same conditions. Scale bar 100um.

**Supplementary Figure 3**. Different AB dilutions were tested in IMC, with representative images showing of ABs against aSMA, podoplanin, CD90 and Tenascin-C on HNSCC tissue. Scale bar 100um.

**Supplementary Figure 4**. Visualization of Ki-67 (red), CD31 (yellow), CD146 (magenta), podoplanin (blue) and EGFR (cyan) using IMC and IHC in the HNSCC cTMA core. Scale bar 100 μm

**Supplementary Figure 5**. Visualization of various immune subsets in HNSCC core. Natural killer (NK) cells (yellow) and cytotoxic T-cell (CD8+, red) can be observed in the tumor epithelia, whereas majority of T-regulatory cells (CD4+, blue), B cell (CD20+ cyan) and M2-like macrophages (CD163+, green) can be observed in the tumor stroma. Scale bar 100 μm. B) and C) Correlation plots of positive cells as detected by IMC and IHC and counted for each marker.

**Supplementary Figure 6**. FSP-1 (red), CD140β (blue), CD90 (yellow), caveolin-1 (green) and YAP-1 (magenta) can be successfully visualized in the osteosarcoma tissue section. Scale bar-100 μm.

**Supplementary Figure 7**. A) and B) Number of cells and relative percentage in each cluster related to tissue type. C) Distribution of cells according to tissue type visualized after dimensionality reduction using t-SNE. D) Distribution of cells based on marker expression and phenotype in the same t-SNE.

