## Supplementary figures and images for "Development of a high dimensional imaging mass cytometry panel to investigate spatial organization of tissue microenvironment in formalin-fixed archival clinical tissues"

### Figure S1

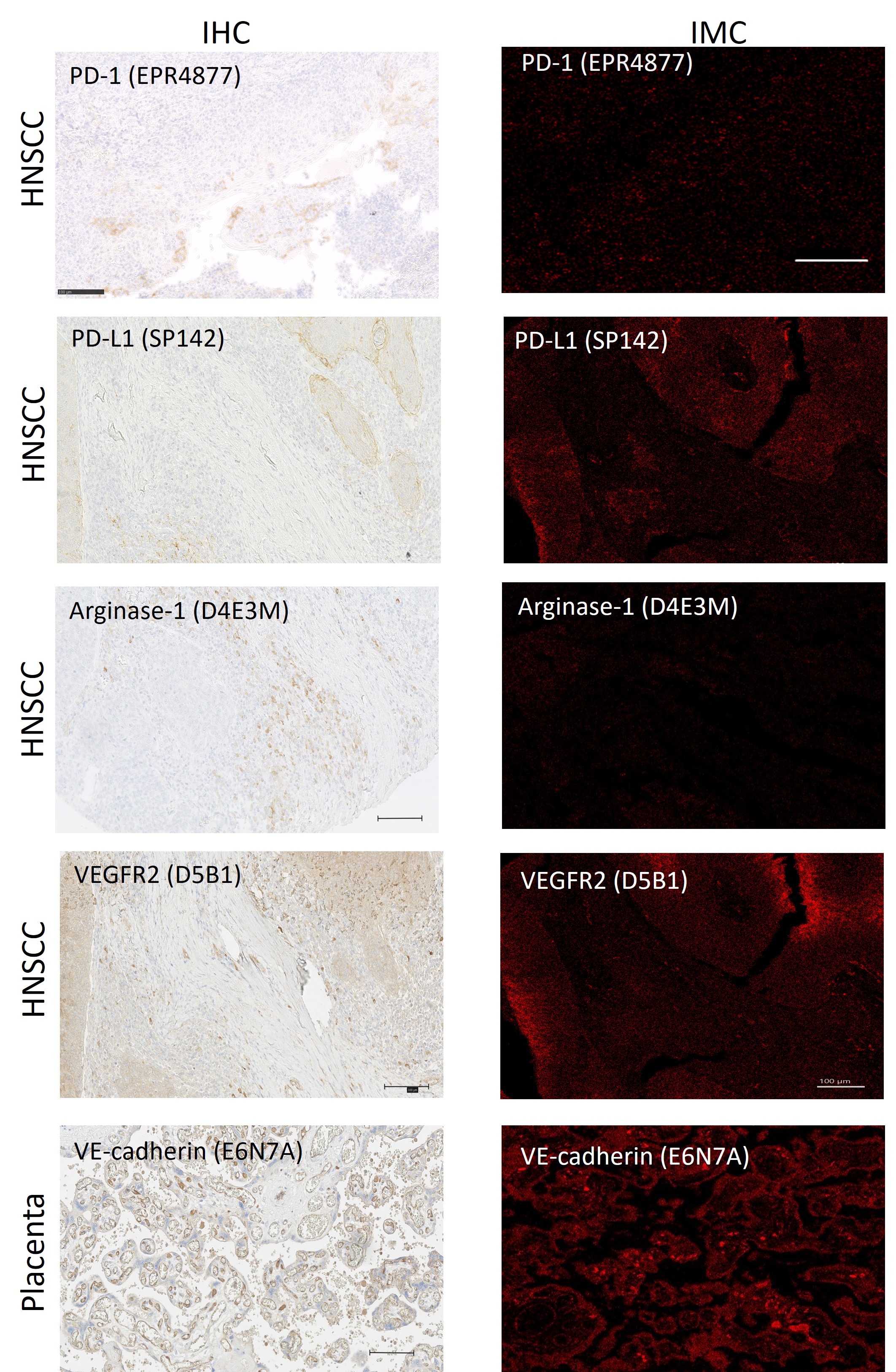

### Figure S2

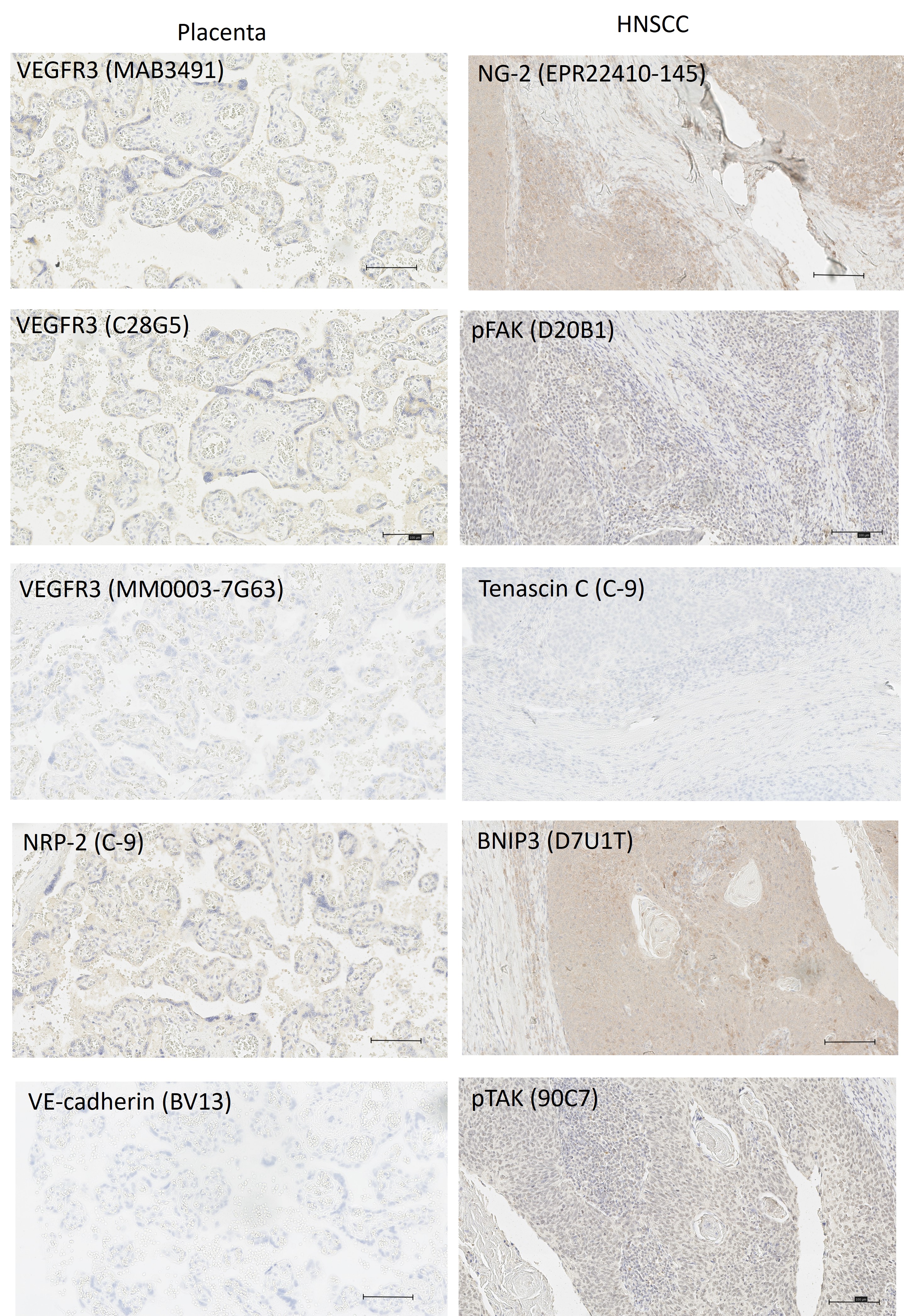

### Figure S3

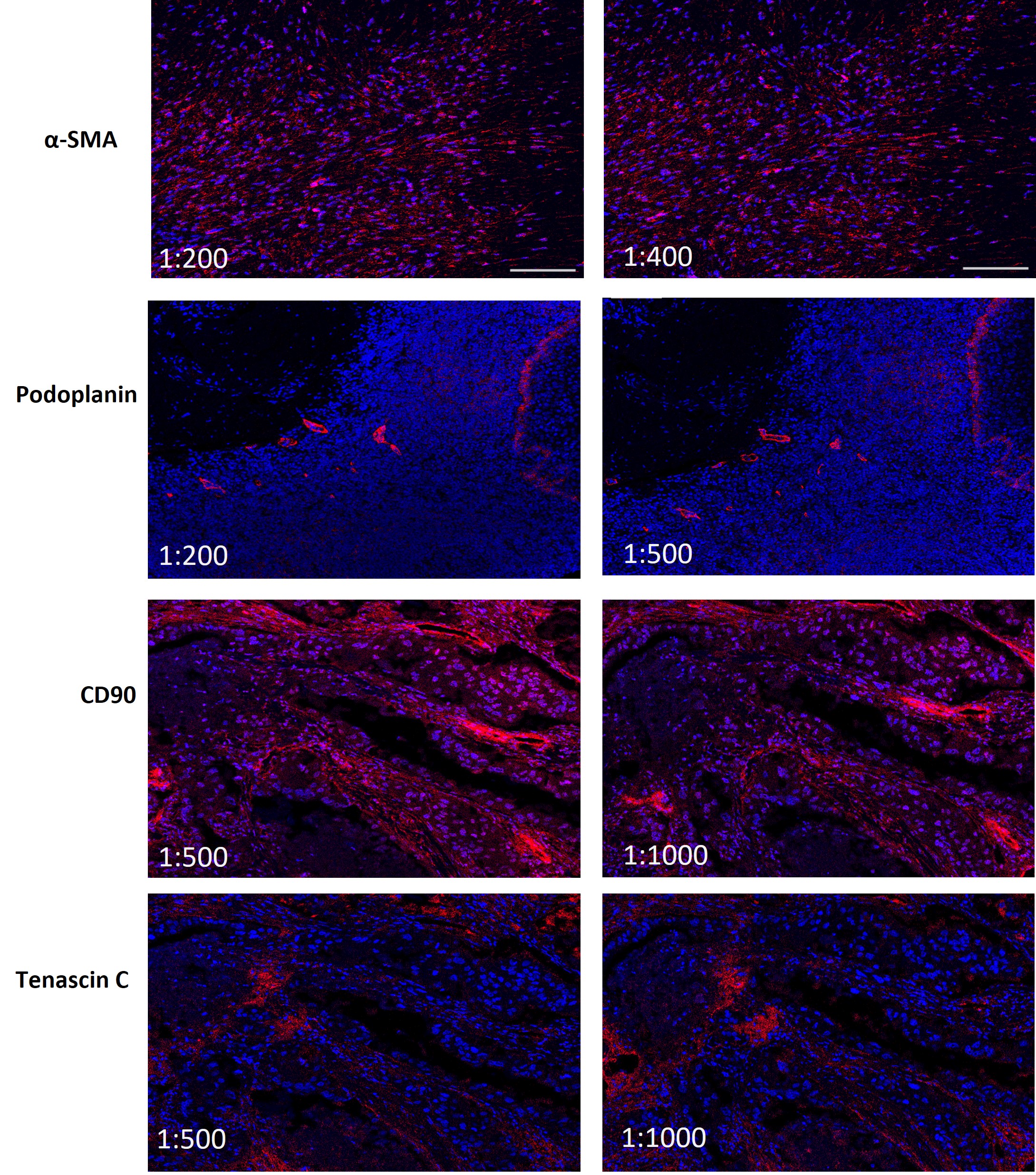

### Figure S4

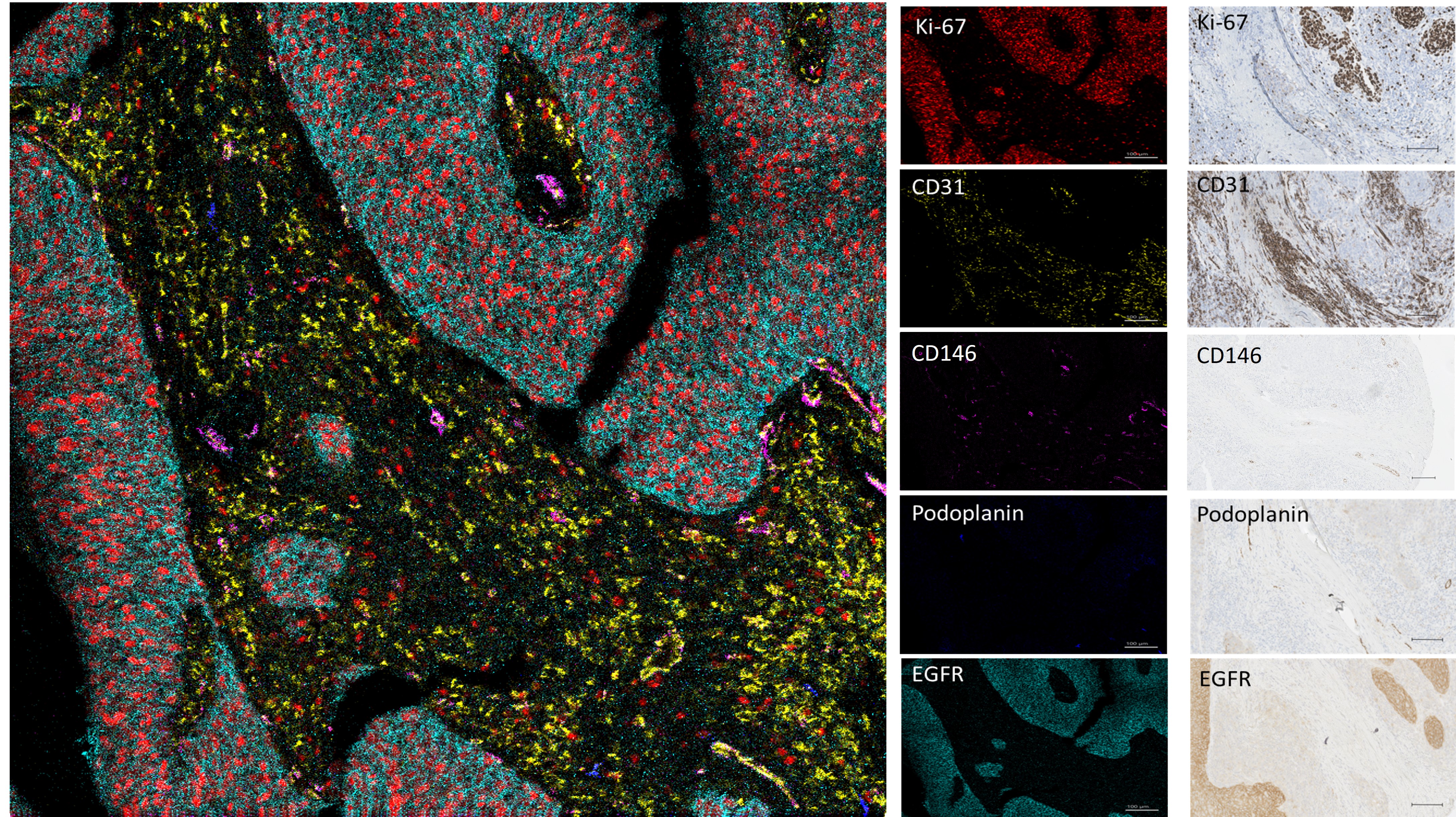

### Figure S5

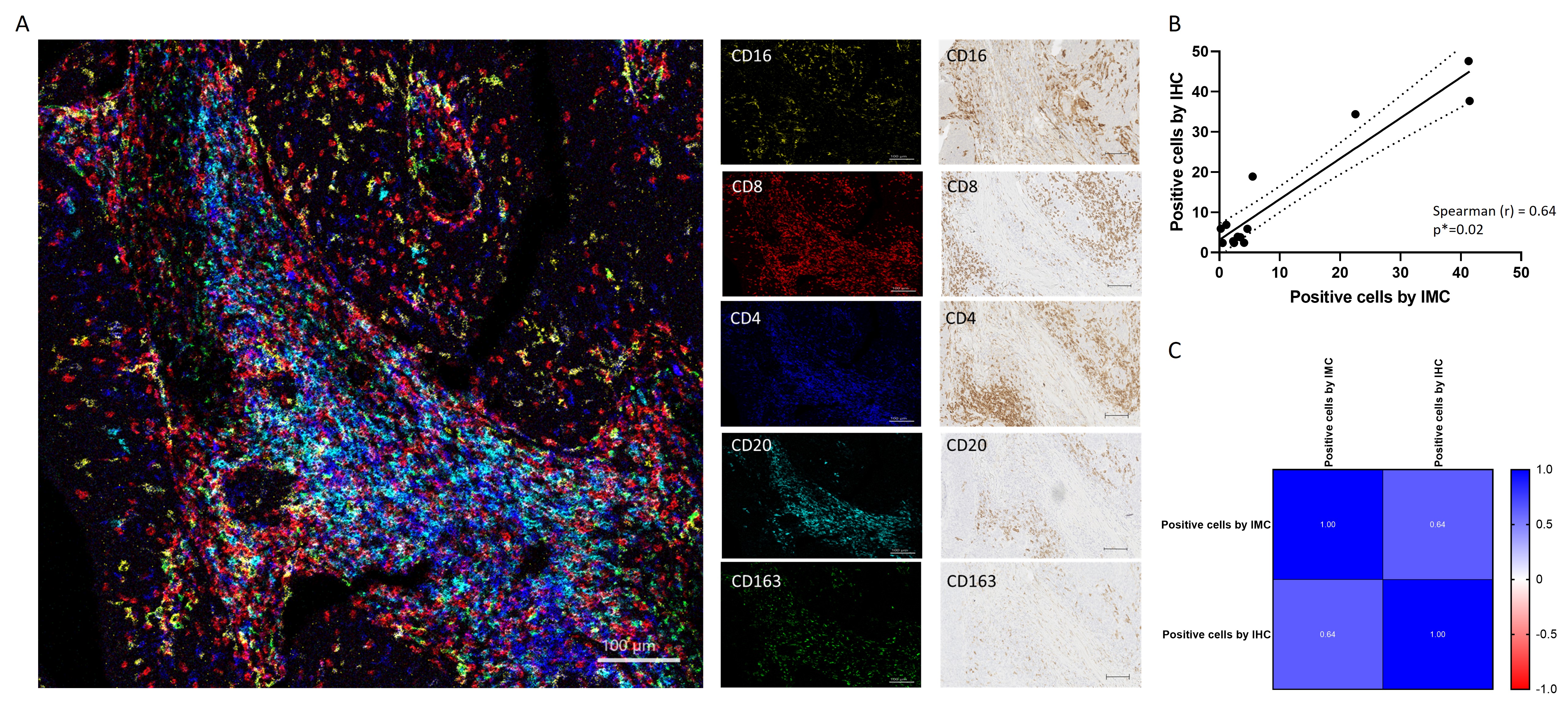

### Figure S6

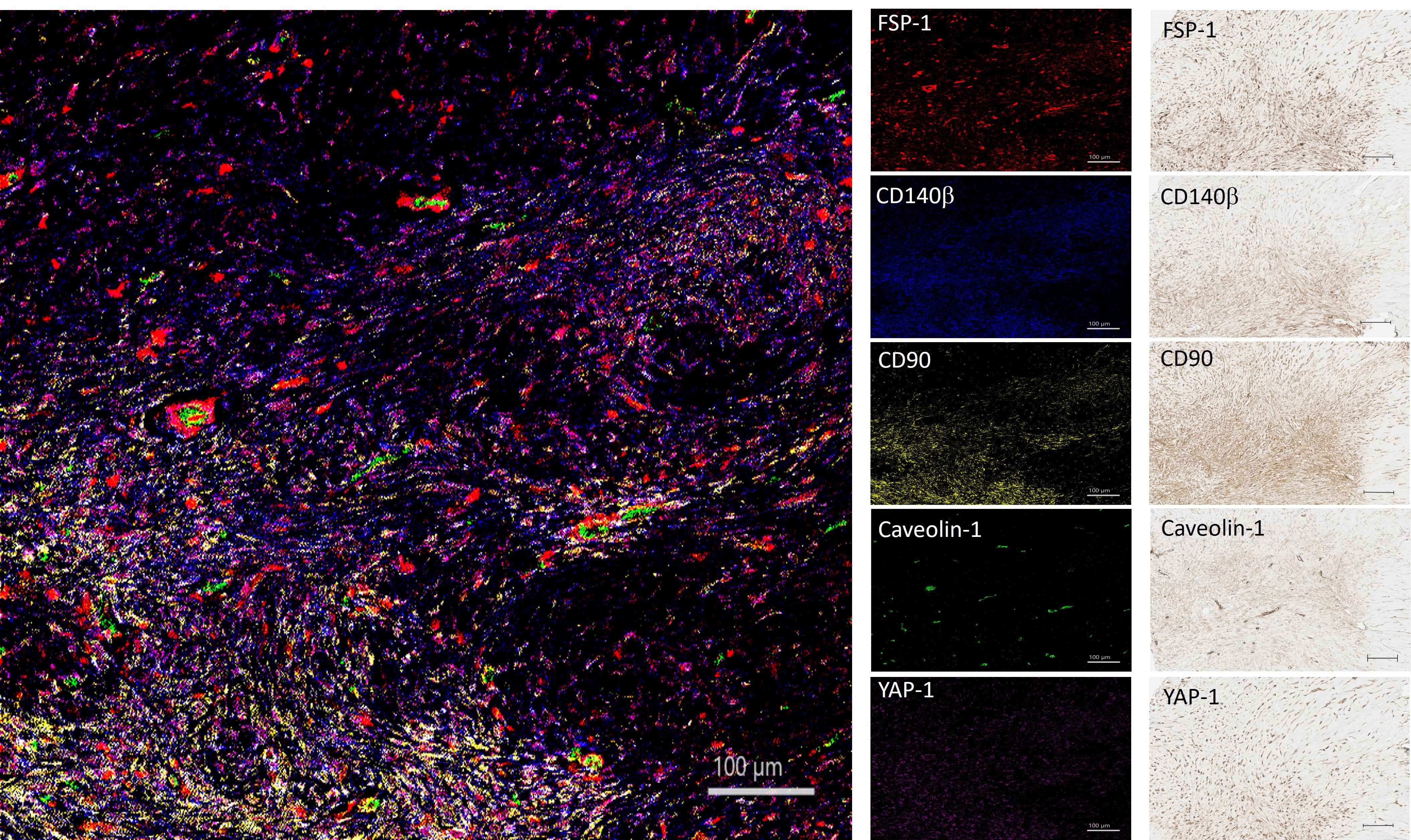

### Figure S7

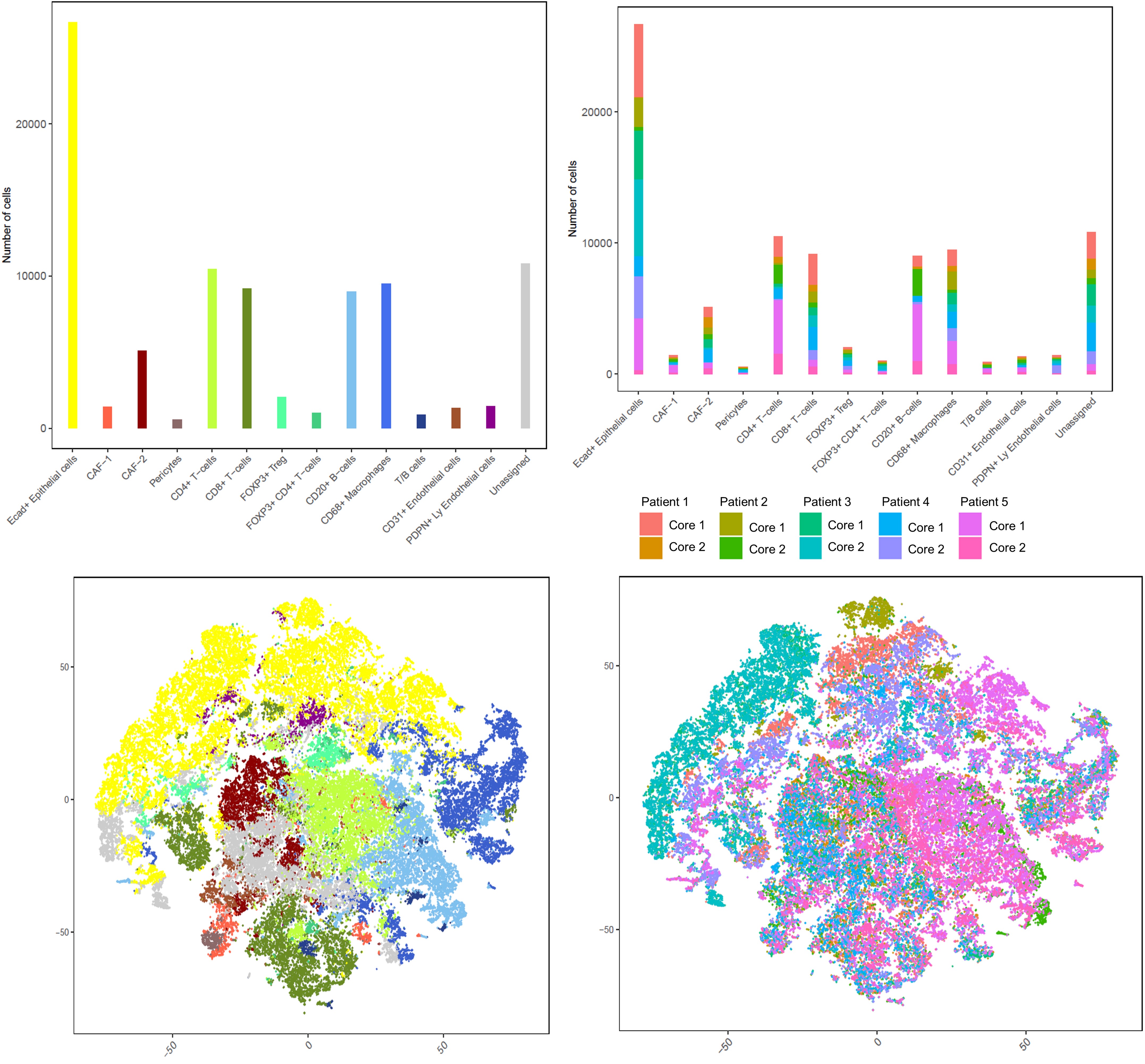

### Supplementary Table 1

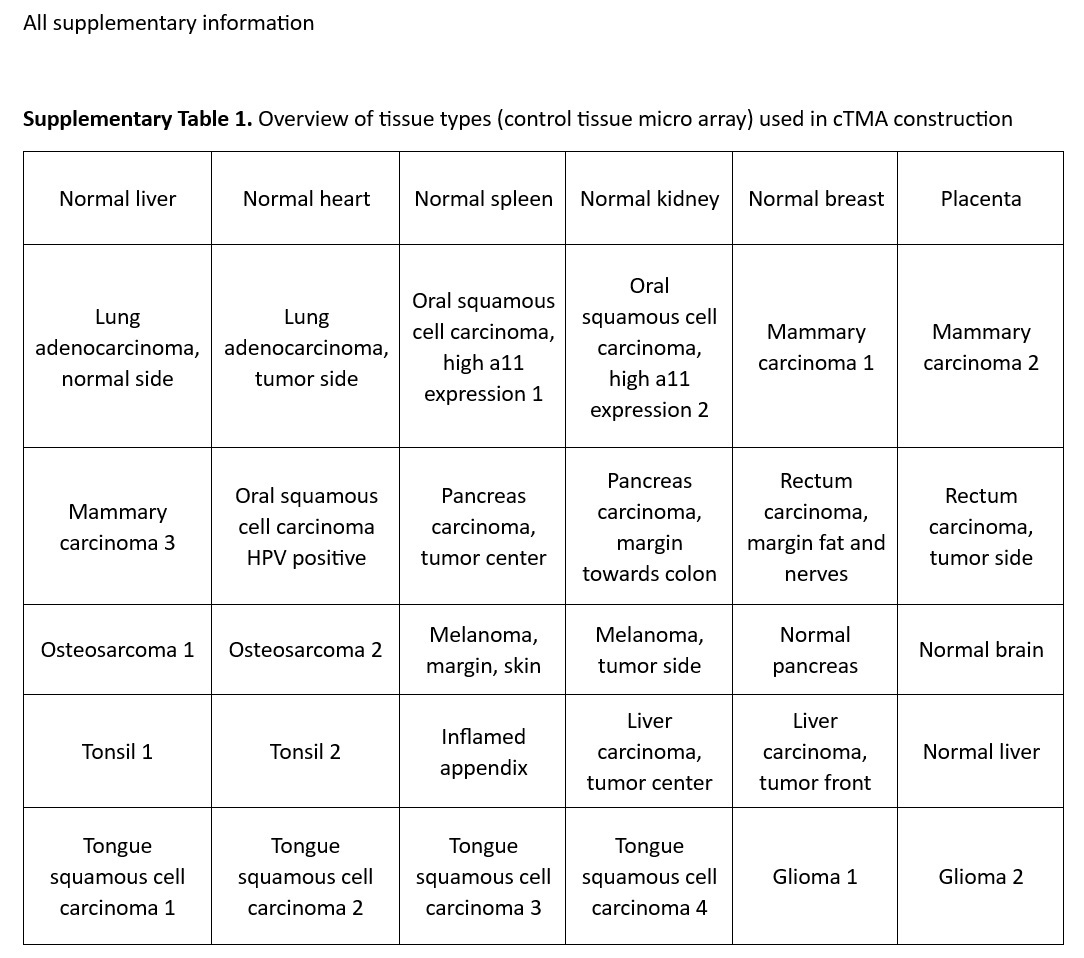

### Supplementary Table 2

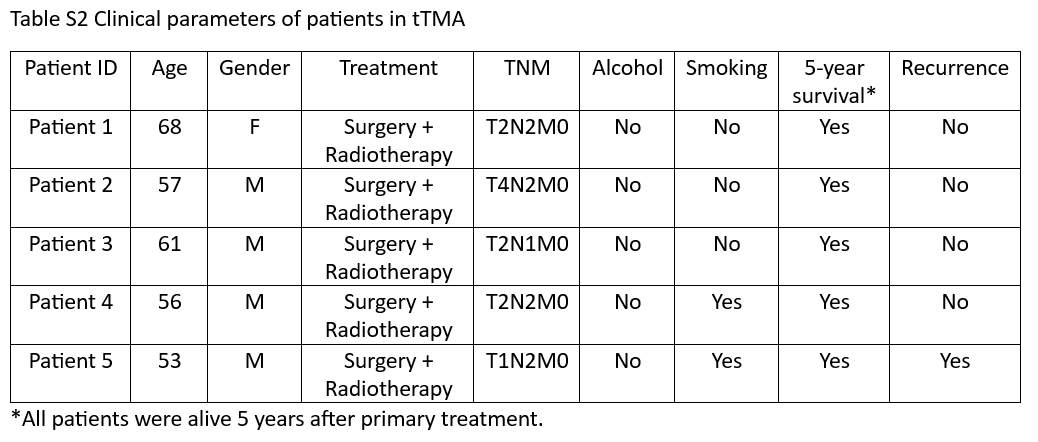
